# AMPK-p38 axis converts human pluripotent stem cells to naïve state

**DOI:** 10.1101/2022.03.31.486536

**Authors:** Zhennan Yang, Yajing Liu, Huaigeng Xu, Junko Yamane, Akitsu Hotta, Wataru Fujibuchi, Jun K Yamashita

## Abstract

Pluripotent stem cells (PSCs) have been reported to exhibit two stages of pluripotency, ‘primed’ and ‘naïve’ states. Typical human PSCs (hPSCs) are in the primed state. Though several methods for conversion from primed to naïve state have been reported, the mechanism of the process is not fully understood. Here, we report that 5’ adenosine monophosphate-activated protein kinase (AMPK) and its downstream p38 is a signaling axis that can induce the naïve conversion of hPSCs with single pathway activation. The simple addition of an AMPK activator, 5-aminoimidazole-4-carboxamide-1-β-D-ribofuranoside (AICAR), or overexpression of a constitutive active form of p38 (CA-p38) alone in primed hPSCs induced naïve hPSCs that satisfied naïve state criteria: differentiation ability to three germ layers and naïve state-specific transcriptional expression, epigenomic resetting, and mitochondrial activity. RNA-seq analysis demonstrated that our AICAR- or CA-p38-induced naïve hPSCs show closely similar gene expression patterns to naïve state human embryonic stem cells (HNES1) derived from human inner cell mass (ICM). This novel and simple naïve conversion method provides new avenues for understanding and elucidating the fundamental mechanism of naïve conversion.

## INTRODUCTION

Pluripotent stem cells (PSCs) have been reported to exhibit two stages of pluripotency, ‘primed’ and ‘naïve’ states. The ‘naïve’ state is closer in characteristics to the inner-cell mass (ICM) of blastocysts in the preimplantation stage, while the primed state is closer to the epiblast in the postimplantation embryo (Brons et al., 2007; Nichols et al., 2009; Boroviak et al., 2014; Boroviak et al., 2015; Weinberger et al., 2016). The primed state cells are considered not capable of chimera formation, while the naïve state cells may be capable of chimera formation (Masaki et al., 2015; Hu et al., 2020). In addition, naïve state pluripotent stem cells express pluripotency genes including naïve state-specific markers, and are in a “ground state” with the most lesser epigenetic modification during the whole developmental process (Smith et al., 2012; Marks et al., 2012; Ficz et al., 2013; Yagi et al., 2017). Human PSCs are in the primed state, whereas mouse embryonic stem cells (ESCs) are principally in the naïve state (Thomson et al., 1998; Choi et al., 2016; Weinberger et al., 2016).

Recently, several groups have reported primed-to-naïve state conversion protocols in human PSCs. However, in these studies, the process of primed-to-naïve state conversion is mediated by either combination of small molecules and cytokines (Gafni et al., 2013; Chan et al., 2013; Theunissen et al., 2014; Qin et al.,2016; Szczerbinska et al., 2019; Hu et al., 2020, see Table S1) or a single compound, HDAC inhibitor, that broadly alters cellular epigenetic status (Guo et al., 2017). These multifactorial conversions to the naïve state make it difficult to explore the mechanisms behind their naïve conversion process.

As a breakthrough for this situation, we recently demonstrated that activating the 5’ adenosine monophosphate-activated protein kinase (AMPK) pathway with the simple addition of a single chemical AMPK activators can convert primed state mouse epiblast stem cells (Epi-SCs) to the naïve state embryonic stem cells (Liu et al., 2021). AMPK is an enzyme that is activated when it senses an increase in the AMP/ATP ratio due to energy deficit caused by exercise or caloric restriction and is responsible for regulating cellular energy homeostasis. AMPK activator treatment (5-aminoimidazole-4-carboxamide-1-β-D-ribofuranoside (AICAR), A769662, or metformin) on mouse Epi-SCs induced an increase in various pluripotent and naïve gene expressions and resulted in the appearance of naïve cells.

Here we report the induction of naïve conversion of human primed PSCs with AMPK activator treatment. The addition of AICAR to human primed PSCs cultured in a naïve maintenance medium (PXGL medium) (Bredenkamp et al., 2019) can induce naïve cell appearance. p38 is a critical downstream of AMPK-elicited naïve conversion, revealing its activation alone similarly induces naïve conversion. RNA seq data confirmed that our induced naïve hPSCs are close to naïve state human ESCs (HNES1) (Guo et al., 2016) derived from human ICM in gene expression. Successful naïve conversion of human PSCs through a single signaling pathway would provide a profound understanding of the pluripotency statuses and open avenues for novel regeneration and rejuvenation strategies.

## RESULTS

### AICAR treatment converts human pluripotent stem cells to naïve state

Recently, We have reported that the treatment with AICAR or the expression of a constitutive active form of p38 (CA-p38) can maintain the naïve state of mouse ESCs and convert mouse EpiSC to naïve mouse ESCs (Liu et al., 2019; Liu et al., 2021). Based on these data, we examined whether the treatment with AICAR or CA-p38 could convert primed human PSCs to the naïve state. As an initial attempt, we added AICAR to human ESCs and human iPSCs (Takahashi et al., 2006; Takahashi et al., 2007).

Whereas naïve mouse ESCs can be fully maintained in vitro by culturing them in two small molecule inhibitors (2i) of kinases (MEK and GSK3) and leukemia inhibitory factor (LIF)(2iL) (Sato et al., 2004; Ying et al., 2008; Wray et al., 2011), but human naïve PSCs have required further modification such as the addition of a Wnt inhibitor, XAV939 (Bredenkamp et al., 2019). Therefore, in this study, we used PXGL medium consist of a mixture of PD0325901 (1 μM), XAV939 (2 μM), Go6983 (2 μM), and human LIF (10 ng/ml) in Ndiff227 medium (Bredenkamp et al., 2019) as a naïve human PSC maintenance medium. Then, we examined the effect of AMPK activation in this culture condition (Figure1A).

**Figure1.**
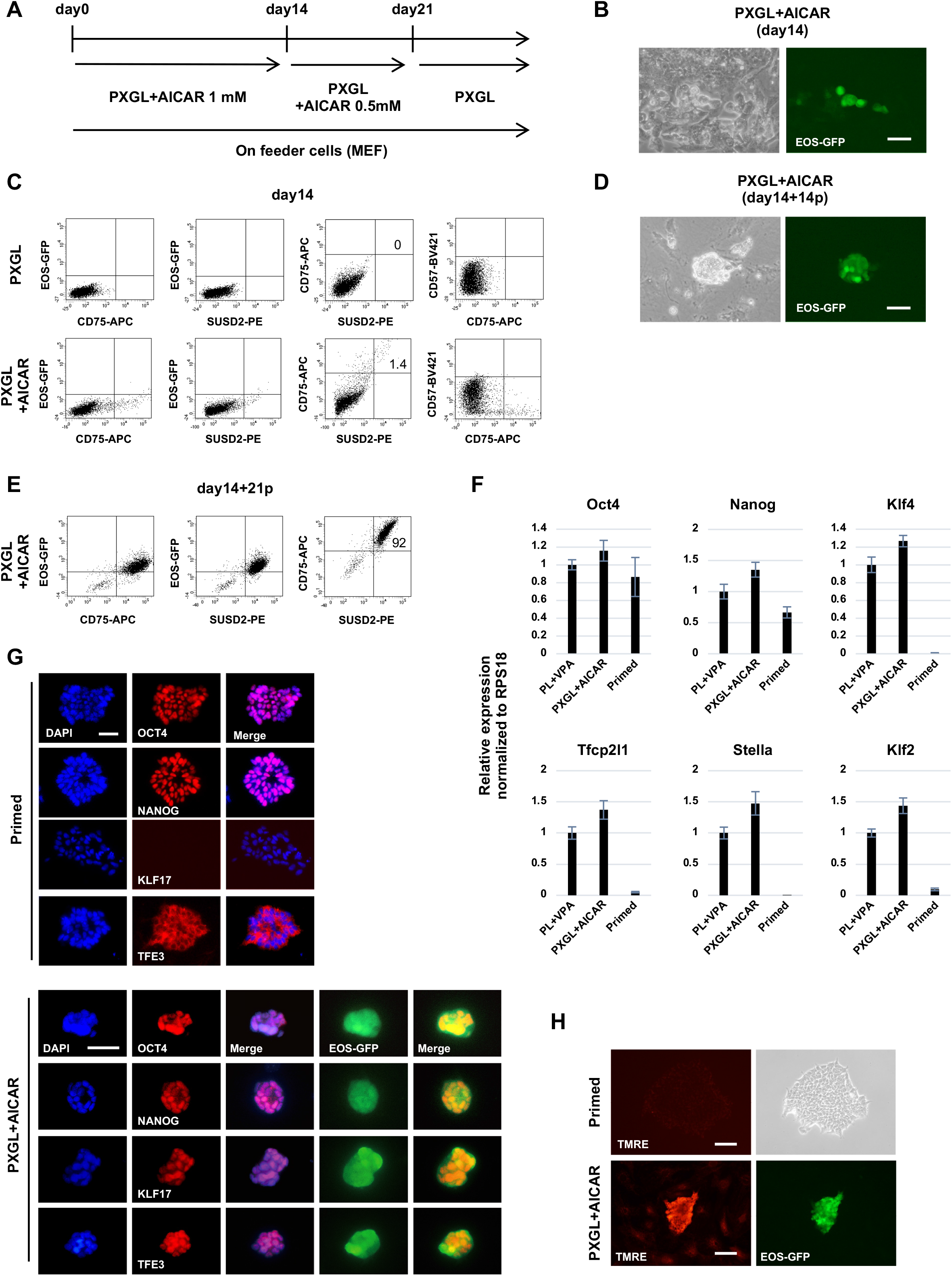
Naive conversion. (A) Naïve conversion protocol using AICAR. PXGL; PD0325901 (1 μM), XAV939 (2 μM), Go6983 (2 μM), and human LIF (10 ng/ml) in Ndiff227 medium. (B) Appearance of EOS-GFP-positive cell clusters induced by AICAR (day14). (C) Flow-cytometry analysis of EOS-GFP, SUSD2, CD75, and CD57 expression (day14). (D) EOS-GFP-positive naïve-like colony after expansion in PXGL (day14+14p). (E) Flow-cytometry analysis of EOS-GFP, SUSD2 and CD75 expression (day14+21p). (F) RT-qPCR analysis of sorted SUSD2+CD75+ cells and parental primed hESCs (H1-EOS). AICAR: day14+13p and VPA: day9+4p. Error bars: S.D. of technical triplicates. (G) Immunostaining for OCT4, NANOG, KLF17, and TFE3 of primed and AICAR-induced cells. AICAR: day14+14p. (H) TMRE staining for mitochondrion dependent on mitochondrial membrane activity. Scale bars: 100 μm in B, D, H; 50 μm in G. AICAR: day14+16p.

The appearance of human naïve PSCs was initially monitored by EOS-GFP reporter expression as a naïve state marker (Takashima et al., 2014, Hotta et al., 2009). We generated human PSCs expressing EOS-GFP reporter gene (see Methods) using H1 human ESCs (H1-EOS) and Ff-I14 human iPSCs (Ff-I14-EOS).

When H1-EOS cells were treated with valproic acid (VPA), which has been reported to induce a primed-to-naïve state transition in human PSCs (Guo et al., 2017), GFP-positive cells were successfully induced (Figure S1A), indicating the validity of the EOS-expressing cell system. When these cells were treated with 1 mM of AICAR in the PXGL medium for 14 days, we observed the appearance of small GFP-positive cell clusters (Figure 1B). Flow cytometry analysis confirmed that very few but distinct appearances of GFP-positive cells were detected, different from cells only treated with PXGL medium. Most cells were negative for CD57, a primed state marker, and some cells started to be positive (less than 2%) for naïve state markers, CD75 and SUSD2 (Collier et al., 2017; Bredenkamp et al., 2019) (Figure 1C). After the emergence of GFP-positive cells (14 days of induction), cells were passaged and cultured for an additional 1 week with a reduced AICAR concentration (0.5 mM) (Figure 1A). Then AICAR was removed, and cells were continued to expand in the naïve maintenance PXGL medium alone on MEF-feeder cells. GFP-positive cells could propagate for over 4 months in the PXGL medium without AICAR to show naïve-like dome-shaped colonies (Figure 1D). Their doubling time was 4-5 days in this culture condition (PXGL with MEF-feeder cells) (Figure S1B). The CD75+/SUSD2+/GFP+ cell population was enriched after passaging and clearly identified by flow cytometry analysis (Figure 1E). RT-qPCR analysis for FACS-purified CD75- and SUSD2-positive naïve-like cells shows that pluripotency markers, Oct4 and Nanog, were expressed in both primed H1-EOS and VPA- and AICAR-induced naïve-like cells. By contrast, naïve state markers, Klf4, Tfcp2l1, Stella, and Klf2 were only expressed in the naïve-like cells (Figure 1F). Immunofluorescence staining also shows the expression of OCT4 and NANOG in both primed state H1-EOS and the naïve-like cells, while naïve state marker KLF17 was only in the naïve-like cells. Nuclear translocation of TFE3 was reported to occur in naïve state cells (Betschinger et al., 2013). Consistent with this, VPA- and AICAR-induced naïve-like cells showed nuclear TFE3 localization, while primed H1-EOS cells did cytoplasmic localization (Figure 1G). TMRE (tetramethyl-rhodamine methyl ester) staining showed an increase of mitochondrial activity, which is a feature of naïve PSCs, in the AICAR-induced naïve-like cells than primed H1-EOS cells (Zhou et al., 2012; Takashima et al., 2014) (Figure 1H). All these results indicate that AICAR-induced cells exhibit various characteristics of the naïve state. In addition, similarly, hiPSC (Ff-I14-EOS) has also successfully induced CD75+/SUSD2+/GFP+ dome-shaped naïve cell-like colonies by AICAR treatment (Figure S1C).

### Differentiation ability of AICAR-induced naïve human pluripotent stem cells

The differentiation potential of AICAR-induced naïve-like cells was evaluated with in vitro differentiation. Since the induction protocols we used were developed for primed state PSCs, AICAR-induced naïve-like cells were once ‘re-primed’ by cultured in a primed cell medium, AK02N, for more than 3 passages. The re-primed cells showed differentiation to three germ layer cell lineages. THY1- or PDGFRβ-positive mesoderm cells appeared after differentiation with a mesoderm induction method, modified DD protocol (Uosaki et al., 2011) (Figure 2A). Endoderm (SOX17- or CXCR4-positive) was induced by treating cells with Activin A and Wnt3A (Kroon et al., 2008) (Figure 2B). Neural differentiation was achieved using Noggin and SB431542 (Chambers et al., 2009) and was confirmed by TUJ1 and MAP2 immunostaining (Figure 2C).

**Figure2.**
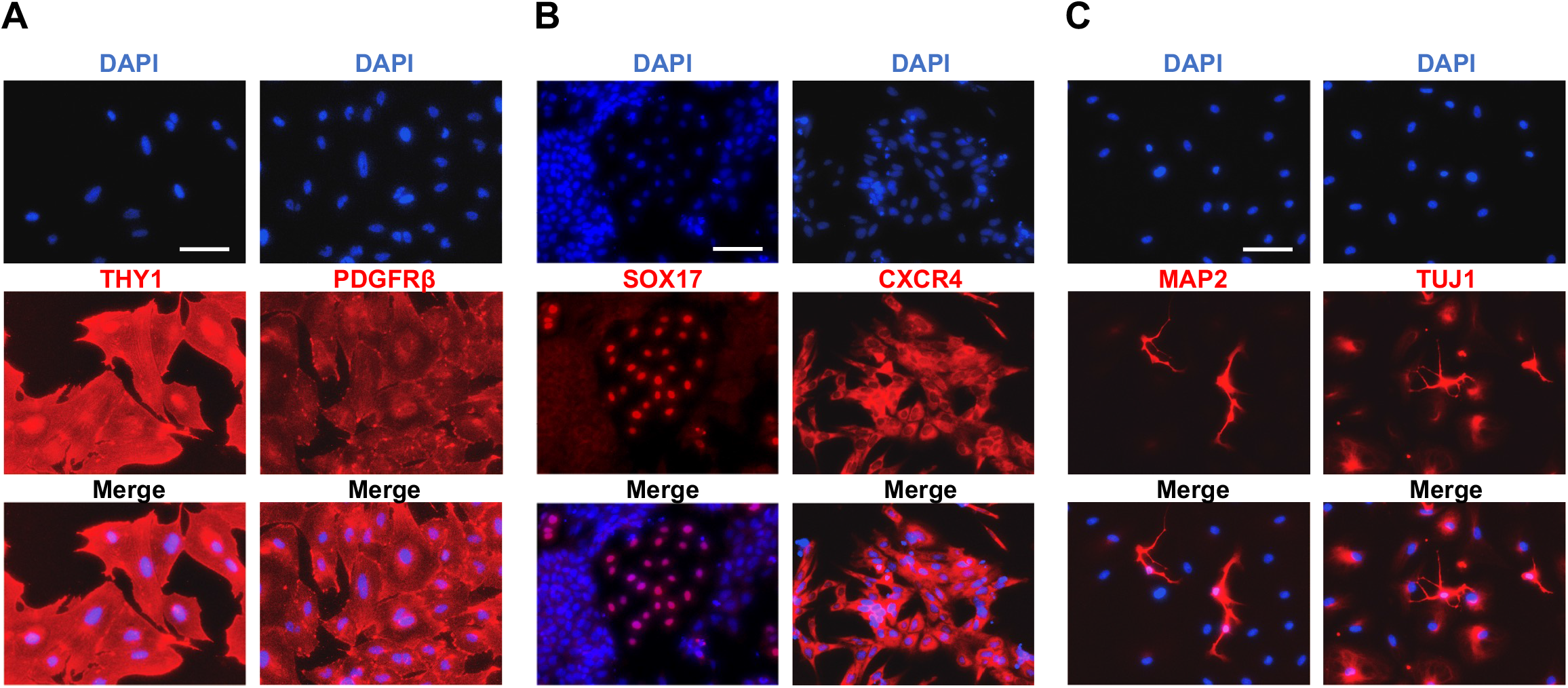
Differentiation of AICAR-induced naïve-like cells. (A) Immunofluorescence for mesoderm markers THY1 and PDGFRβ. (B) Immunofluorescence for endoderm markers SOX17 and CXCR4. (C) Immunofluorescence for ectoderm markers MAP2 and TUJ1. Scale bars: 100 μm.

### Epigenome status

Histone 3 lysine 9 trimethylation (H3K9me3) is one of the heterochromatin markers. Foci formation in H3K9me3 staining is observed in the primed state PSCs while it disappeared in the naïve state (Takashima et al., 2014). Consistent with this, H3K9me3 foci were observed in primed H1-EOS but not in AICAR-induced naïve-like cells (Figure 3A). Human ICM cells, naïve mouse ES cells, and naïve human PSCs exhibit global DNA hypomethylation, whereas hypermethylation is observed in primed mouse Epi-SCs and primed human PSCs (Ficz et al., 2013; Habibi et al., 2013: Leitch et al., 2013; Takashima et al., 2014). We performed immunofluorescent staining of DNA methylation markers, 5-methylcytosine (5mC) and 5-hydroxymethylcytosine (5hmC), and confirmed their expressions were apparently reduced in AICAR-induced naïve-like cells (Figure 3B).

**Figure3.**
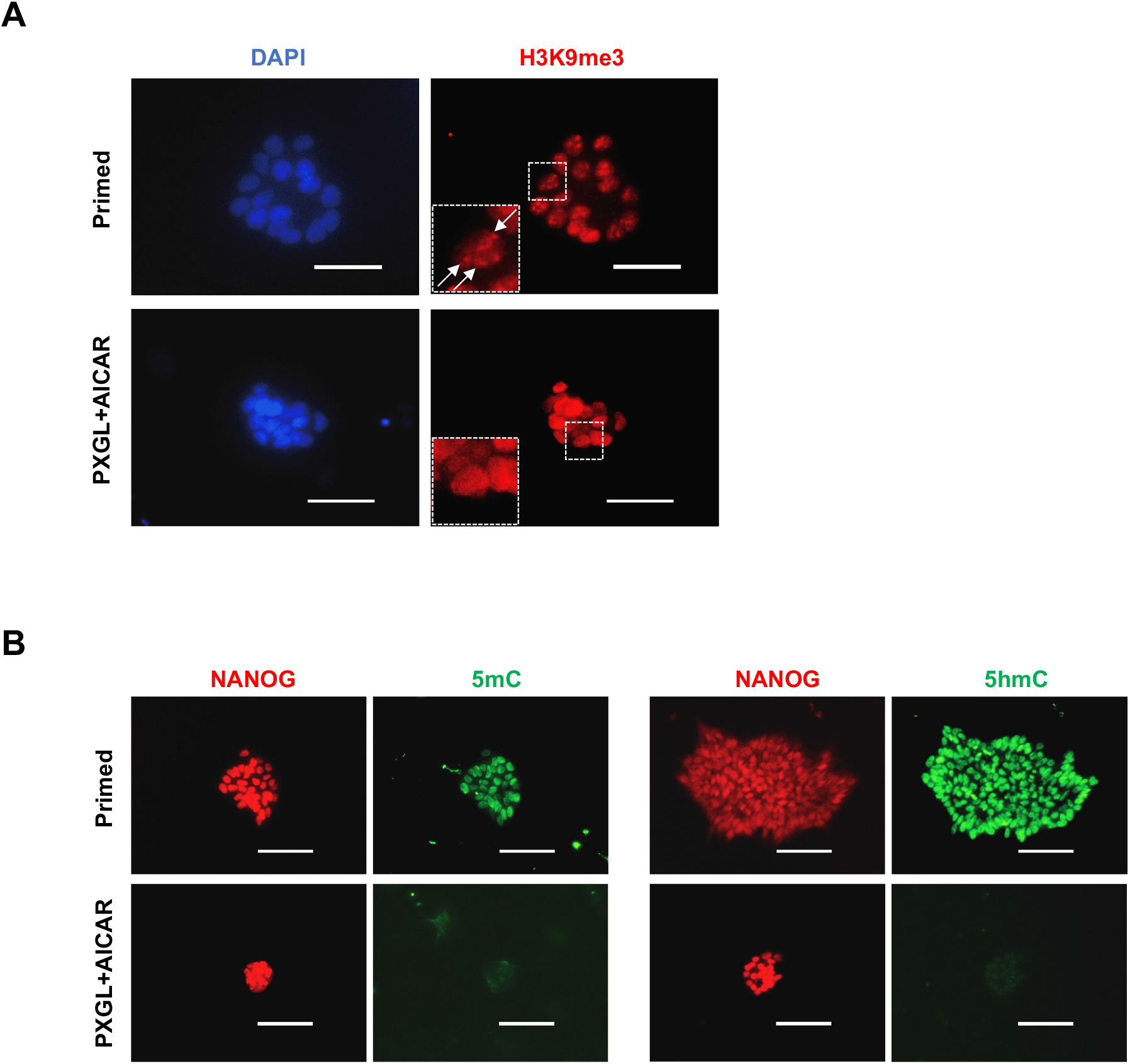
Epigenome analysis of induced naïve hPSCs. (A) Immunostaining for H3K9me3. (B) Immunostaining for 5mC, 5hmC, and NANOG. Scale bars: 50 μm in A; 100 μm in B. AICAR: day14+14p.

All these results confirm that the addition of AICAR alone to PXGL maintenance conditions can induce the conversion of primed human PSCs to cells that satisfy multiple naïve state criteria.

### p38 is an AICAR downstream and can convert human pluripotent stem cells to naïve state

AICAR treatment can influence many downstream signaling pathways involved in broad biological phenomena including metabolism, cell growth, autophagy, and others (Mihaylova et al., 2011; Zang et al., 2009; Li et al., 2005).

We have previously reported that p38 is a critical downstream of AMPK in the maintenance and induction of the naïve state of mouse PSCs (Liu et al., 2019; Liu et al., 2021). Then, we examined the involvement of p38 in the AICAR-elicited naïve conversion of human PSCs. First, the addition of SB203580, a p38 inhibitor, together with AICAR inhibited the AICAR-elicited naïve cell induction, indicating the involvement of p38 in the naïve conversion (Figure 4A).

**Figure 4.**
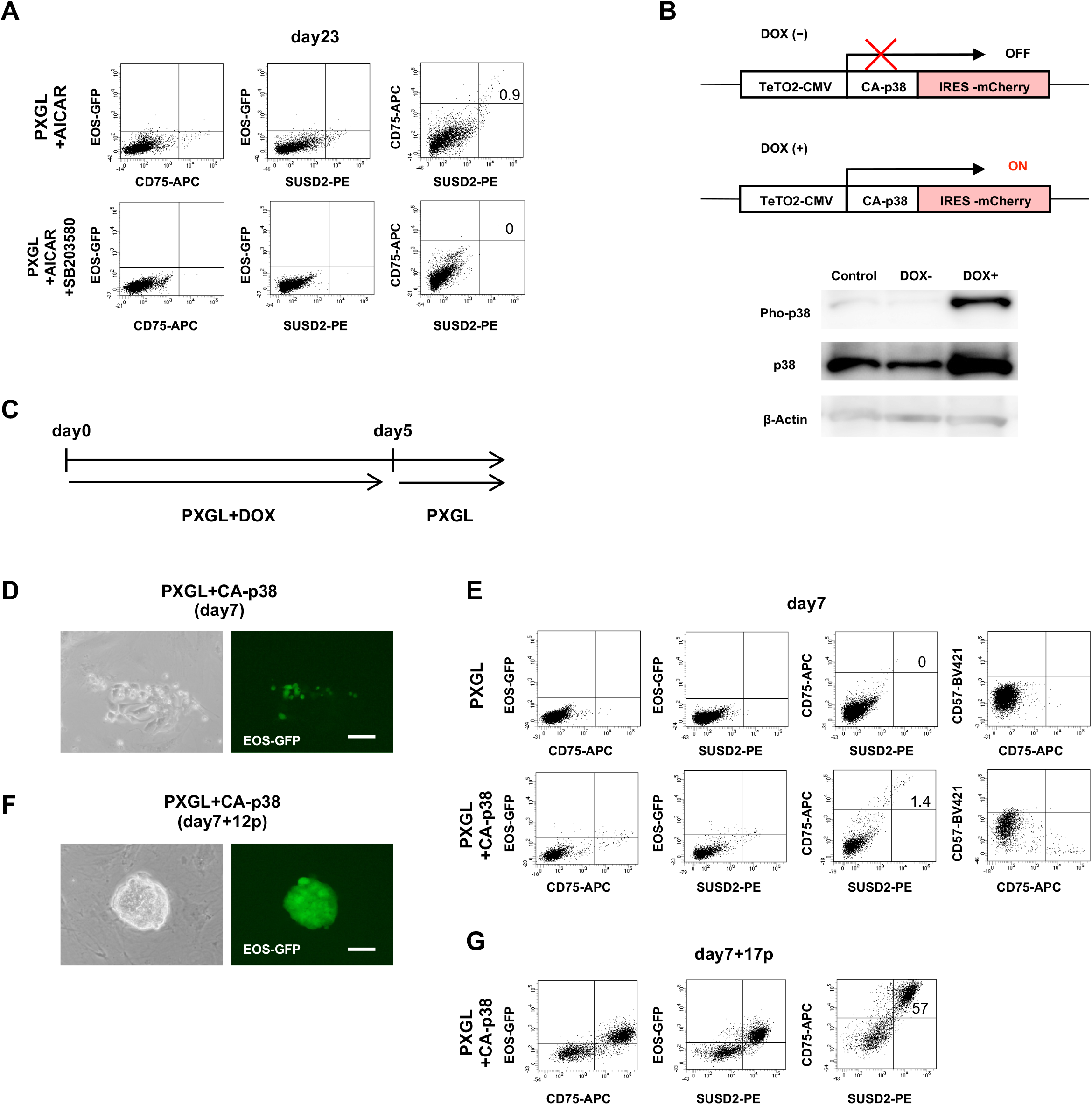
Involvement of p38 in AMPK-mediated naïve conversion. (A) Flow-cytometry analysis of EOS-GFP, SUSD2 and CD75 expression under treatment of p38 inhibitor (SB203580) (day23). (B) Tet-inducible CA-p38 expression system (upper panel). Western blotting for p38 proteins after Dox treatment. β-Actin: loading control. (C) Naïve conversion protocol with p38 activation. (D) Appearance of EOS-GFP-positive cell cluster induced by CA-p38 (day7). (E) Flow-cytometry analysis of EOS-GFP, SUSD2, CD75, and CD57 expression after induction of CA-p38 (day7). (F) EOS-GFP-positive naïve-like colony after expansion in PXGL (day7+12p). (G) Flow-cytometry analysis of EOS-GFP, SUSD2 and CD75 expression after expansion in PXGL (day7+17p). Scale bars: 100 μm.

Next, we verified whether p38 could induce naïve state conversion. We generated human PSCs in which p38 pathway can be activated by the drug-inducible expression of constitutive active form p38 (CA-p38). We transfected a tetracycline-inducible (Tet-ON) CA-p38 into H1-EOS (CA-p38-H1-EOS) cells (Xu et al., 2013; Liu et al., 2019). Thus, p38 can be phosphorylated and activated in these cells with doxycycline (DOX) treatment (Figure 4B, Figure S2A). When p38 was activated with DOX treatment for 5 days in PXGL medium (Figure 4C), EOS-GFP-positive cell clusters were started to be observed (Figure 4D). Flow cytometry analysis showed the appearance of CD75+/SUSD2+/CD57-/GFP^dul1^ cells (less than 2%), similar to that with AICAR treatment (Figure 4E). After that, CA-p38-induced GFP-positive cells were cultured and propagated in the PXGL medium on MEF feeder cells without DOX. CA-p38 induced cells were maintained for over 4 months with a doubling time of about 4 days (Figure S2B). After the propagation, CA-p38-induced GFP-positive cells formed dome-shaped naïve cell-like colonies (Figure 4F), and were positive for EOS-GFP, CD75, and SUSD2 in flow cytometry (Figure 4G). CA-p38-induced CD75 and SUSD2 double-positive cells were sorted by FACS and analyzed with RT-qPCR. These cells expressed comparable pluripotency markers, Oct4, Nanog, and higher naïve state markers, Klf4, Tfcp2l1, Stella, and Klf2 than parental primed cells (H1-EOS-GFP) (Figure S2C). These CA-p38-induced cells showed pluripotency and naïve state marker protein expressions as well as TFE3 nuclear localization (Figure S2D). The CA-p38 induced GFP-positive cells also exhibited mitochondrial activation with TMRE staining (Figure S2E) and naïve state characteristics in their epigenome (Figure S3), similar to those of AICAR-induced naïve cells. In addition, we confirmed that the re-primed CA-p38-induced cells have the potential for differentiation into all three embryonic germ layers (Figure S4). From these data, we confirmed the success in naïve conversion of hESCs by CA-p38 activation.

### Global gene Expression Analysis of primed and naïve state hPSCs

Finally, we confirmed the naïve state cell induction with AICAR and CA-p38 using global gene expression profiles of RNA-seq analysis. The previously published analysis data of genuine naïve state HNES1 (Guo et al., 2016) derived from human ICM, naïve hPSCs converted from primed cells with various methods (Takashima et al., 2014; Theunissen et al., 2014; Sperber et al., 2015; Guo et al., 2017; Yang et al., 2017), and each primed state cells of them were obtained (see Method). We collected RNA-seq data of naïve cells induced with AICAR, CA-p38, and VPA and each parent primed cells and compared all these samples. Principal component analysis (PCA) revealed HNES1 and chemically reset (VPA) cells both established in the same laboratory (Guo et al., 2016; Guo et al., 2017) are very closely placed. Naïve cells induced in this study (induced with AICAR, CA-p38, and VPA) and 5iL/A-induced naïve cells (Theunissen et al., 2014) form a dense cluster near the cluster with HNES1 and cR cells (Figure 5A). EPS cells (Yang et al., 2017) and NHSM/4i-induced cells (Gafni et al., 2013), previously reported as a kind of naïve cells, however, they are placed near primed cells. Heatmap analysis for 66 genes related to naïve and primed pluripotency (Takashima et al., 2014; Theunissen et al., 2014; Guo et al., 2016; Guo et al., 2017; Hu et al., 2020) further clearly confirmed that AICAR- and CA-p38-induced naïve cells showed similar gene expressions to those of other naïve cells including HNES1 cells (Figure 5B). These data again revealed induced naïve hPSCs such as chemically reset (VPA) cells (Guo et al., 2017), reset cells (Takashima et al., 2014), 5i/L/A-induced naïve cells (Theunissen et al., 2014), and our AICAR- and CA-p38-induced as well as VPA-induced naïve cells form a cluster distinct from each parental primed cells (Figure 5B). From all these data, we concluded that AICAR or p38 activation could induce naïve conversion in human primed PSCs.

**Figure5.**
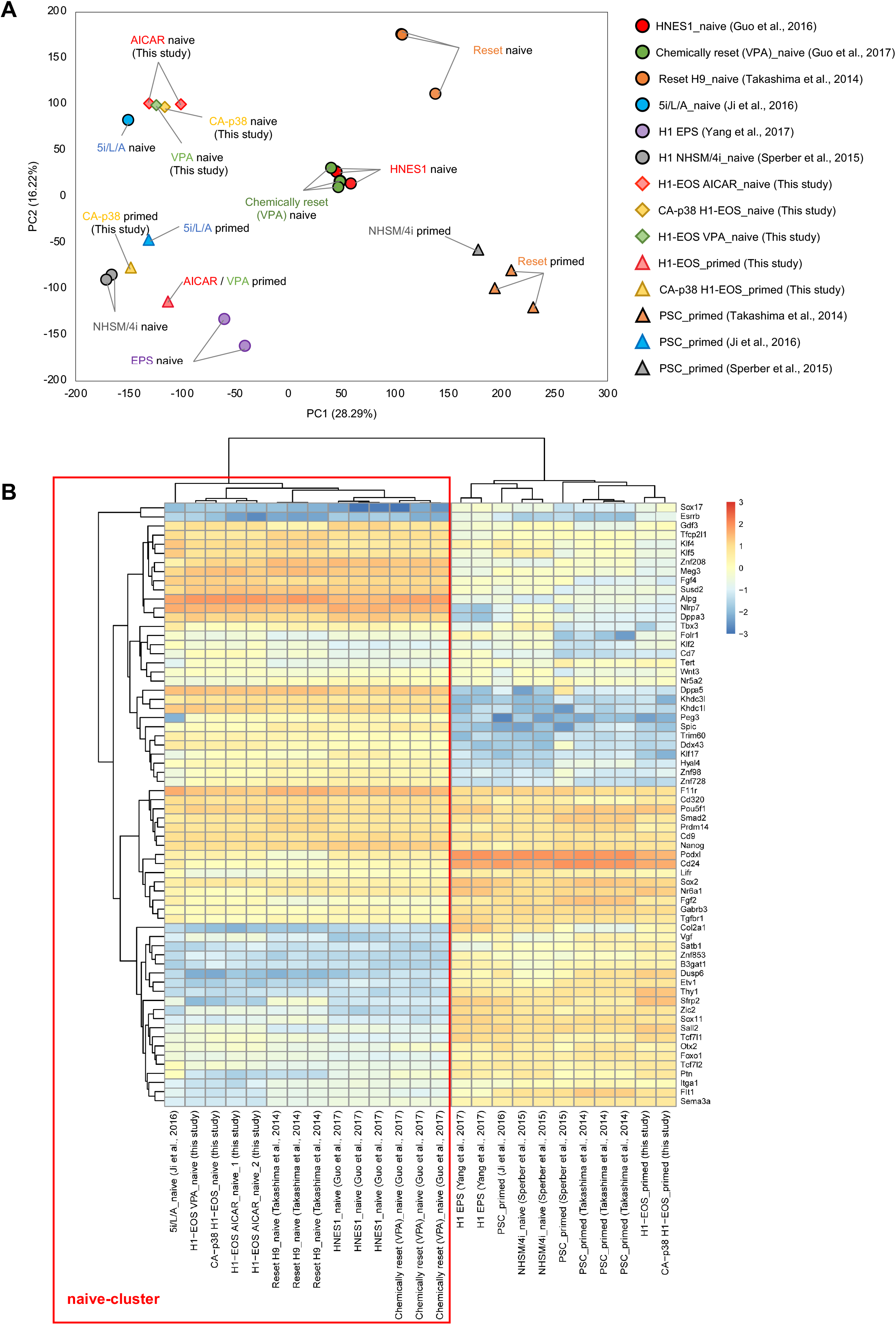
Comparative Expression Analysis of primed and naive state hPSCs. (A) PCA analysis for various naïve and primed hPSCs. (B) Heatmap for RNA expression of selected naïve and primed gene set. Red box: naïve cell cluster.

## DISCUSSION

In this study, we showed that the activation of the AMPK-p38 axis has the potential to convert primed human PSCs to the naïve state in the PXGL conditions. These naïve state PSCs induced by activating the AMPK-p38 axis show the typical naïve state features regarding their marker gene expressions, epigenome status, mitochondrial activity, and global transcriptional expression.

We recently demonstrated that the AMPK-p38 axis is a pathway involved in the maintenance of the naïve state of mouse ESCs (Liu et al., 2019), and it can convert primed mouse Epi-SCs to the naïve state (Liu et al., 2021). Though the core signaling that induces naïve conversion is the same between mouse and human cells, several culture conditions are different (Martin et al., 1981; Bredenkamp et al., 2019). Unlike the naïve conversion of mouse Epi-SCs, Ndiff227 medium was suitable for the naïve conversion of human PSCs in this study. On the other hand, whereas the existence of serum was critical in the mouse naïve conversion with AMPK-p38, the PXGL medium with serum was not suitable for human cells (data not shown). Some differences and conflicts in the naïve conversion conditions are also observed. Wnt activators, GSK3 inhibitors such as CHIR99021, are widely used to induce the naïve conversion of human PSCs in other reports. The recently reported PXGL medium (Bredenkamp et al., 2019), in which XAV939, a canonical Wnt inhibitor, was used instead of GSK3 inhibitors, was suitable in our condition. PXGL medium alone never induced naïve conversion but was suitable for a basal medium for the naïve conversion with AMPK activation and maintenance and expansion of the converted human naïve PSCs. Whereas MEF feeder cells were commonly essential during the naïve conversion process in mouse and human cells, feeder-free culture could maintain mouse naïve PSCs with 2iL medium and human naïve PSCs in PXGL medium, respectively, after the naïve conversion has been completed. Other various differences are also observed. A naïve conversion of hESCs (3iL) using BMP and AMPK inhibitor Dorsomorphin in the TeSR medium supplemented with 2i (MEK and GSK3 inhibitor) and hLIF was reported (Chan et al., 2013). As dorsomorphin is a broad inhibitor for BMP and AMPK pathways, it is still unclear whether the effect was actually through AMPK inhibition. In addition, we failed to induce naïve conversion with AICAR using mTeSR medium instead of Ndiff227 medium (data not shown), suggesting that the combination of medium and compounds may affect the results. The first report of the naïve conversion method (NHSM) (Gafni et al., 2013) used the p38 inhibitor SB203580 as an essential component for naïve conversion, which is the opposite of our study. RNA-seq analysis (Figure 5) reveals that our cells exhibit naïve state PSC characteristics similar to those in reset-cells (Takashima et al., 2014) and chemically reset-cells (Guo et al., 2017), which transcriptional expression is closer to genuine HNES from human ICM (Guo et al., 2016). The naïve cells generated by NHSM method (NHSM/4i_Naïve) (Gafni et al., 2013) belong to another cluster together with various primed cells (Figure 5B), suggesting that p38 activation but not inhibition would be more suitable for the naïve conversion. Recently, Hu et al. reported briefly inhibiting mTOR with Torin1 converted hPSCs to a naïve state (Hu et al., 2020). Since mTOR inhibition is one of the downstream effects of AMPK activation (Agarwal et al., 2015), we also tried inhibiting mTOR with the widely used inhibitor rapamycin in our culture conditions, but we failed to induce the naïve state (data not shown). In addition, as critical human naïve pluripotency markers, such as Khdc1l, Klf4, Klf5, Klf17, Dppa3, Tfcp2l1, Dppa5, and Gdf3 were not significantly upregulated in their naïve hPSCs (Hu et al., 2020) but upregulated in our naïve cells (Figure 5B), we speculate that their naïve PSCs may have a different character to ours. Furthermore, since CA-p38 over-expression alone in the PXGL medium converted our cells to the naïve state (Figure 4), we believe p38 would be one of the critical downstream targets of AMPK in the naïve conversion effect.

Most of the previous reports for naïve conversion of human PSCs used combinations of multiple gene expressions, growth factors, or chemicals in addition to basal naïve maintenance conditions such as 2i (MEK and GSK3 inhibitor) and hLIF. VPA an HDAC inhibitor, has been reported as a robust naïve conversion inducer with a single component (Guo et al., 2017). However, VPA broadly affects gene expressions, so that it would still be required to investigate specific signaling pathways for the naïve conversion mechanism. This study demonstrated that the activation of the AMPK-p38 axis alone could evoke the naïve conversion (Figure 4E), largely contributing to narrowing down the signaling pathway for naïve conversion. In addition, they might potentially lead to applications such as the generation of a stem cell-based embryo model (Yanagida et al., 2021), the induction of primitive endoderm (Guo et al., 2021), or the study of interspecies chimera formation (Kobayashi et al., 2010; Masaki et al., 2015). Thus, this novel and simple naïve conversion method would contribute to profoundly understand the pluripotency statuses and open avenues for novel regeneration and rejuvenation strategies.

## ACKNOWLEDGMENTS

We are grateful to Dr. Yasuhiro Takashima (CiRA: Center for iPS Cell Research and Application, Kyoto, Japan) for valuable advices and discussion, and thank to Xinxiu Xu (The National Center for Drug Screening, Shanghai, China) for providing the plasmid of p38 constitutive activation. This work was supported by AMED grants from Core Center for iPS Cell Research (JP21bm0104001) and Grant-in-Aid for Scientific Research on Innovative Areas (17H19676, 19H05563), Ministry of Education, Culture, Sports, Science and Technology (MEXT), Japan.

## AUTHOR CONTRIBUTIONS

Z.Y. and J.K.Y. conceived the approach. Z.Y. designed, performed, and performed cell culture experiments with contribution of Y.L. A.H. and H.X contributed to EOS reporter experiments. J.Y. helped the RNA-seq experiments. W.F carried out computational work of RNA-seq. Z.Y. and J.K.Y. wrote the paper. J.K.Y. supervised the study and was responsible for funding.

## KEY RESOURCES TABLE

| REAGENT or RESOURCE | COMPANY | Cat No |
| --- | --- | --- |
| **Antibodies** |  |  |
| Mouse anti-SUSD2 Antibody conjugated to PE (clone W5C5) | Bio Legend | Cat# 327406, RRID:AB_940654 |
| Mouse anti-CD75 Antibody conjugated to Alexa 647 (clone ZB55) | BD Biosciences | Cat# 566350, RRID:AB_2869751 |
| Mouse anti-CD57 Antibody conjugated to BV421 (clone ZB55) | BD Biosciences | Cat# 563896, RRID:AB_2632391 |
| Mouse anti-Oct3-4 Antibody (clone C10) | Santa Cruz Biotechnology | Cat# sc-5279 |
| Rabbit anti-NANOG Antibody (clone D73G4) | Cell Signaling Technology | Cat# 4903 |
| Mouse anti-NANOG Antibody (clone 23D2-3C6) | Thermo Fisher Scientific | Cat# MA1-017 |
| Rabbit anti-KLF17 | Sigma | Cat# HPA024629 |
| Rabbit anti-TFE3 | Sigma | Cat# HPA023881 |
| Rabbit anti-Thy1 | Gene Tex | Cat# GTX130072 |
| Mouse anti-PDGFR-β (clone PR7212) | R&D Systems | Cat# MAB1263 |
| Goat anti-SOX17 | R&D Systems | Cat# AF1924 |
| Mouse anti-CXCR4 Antibody conjugated to PE (clone 12G5) | Bio Legend | Cat# 306506 |
| Rabbit anti-MAP2 | Cell Signaling Technology | Cat# 4542 |
| Mouse anti-TUJ1 (clone 2G10) | Sigma | Cat# T8578 |
| Rabbit anti-H3K9me3 | Active Motif | Cat# 39765 |
| Mouse anti-5mC (clone 33D3) | Active Motif | Cat# 39649 |
| Rabbit anti-5hmC | Active Motif | Cat# 39769 |
| Mouse-GFP (clone 3E6) | Thermo Fisher Scientific | Cat# A-11120 |
| **Chemicals, Peptides, and Recombinant Protein** |  |  |
| Recombinant human LIF | FUJIFILM Wako | Cat# 121-06663 |
| Recombinant human LIF | Gibco | Cat# PHC9481 |
| Recombinant human Activin A | R&D Systems | Cat# 338-AC-010 |
| Recombinant human bFGF | FUJIFILM Wako | Cat# 068-04544 |
| Recombinant human Noggin | R&D Systems | Cat# 6057-NG |
| Recombinant human Wnt-3a | Proteintech | Cat# HZ-1296 |
| AICAR | FUJIFILM Wako | Cat# 011-22533 |
| Valproic acid sodium salt | Sigma | Cat# P4543 |
| PD0325901 | Sigma | Cat# PZ0162 |
| XAV939 | Millipore | Cat# 575545 |
| Gö6983 | FUJIFILM Wako | Cat# 078-06441 |
| Y-27632 | FUJIFILM Wako | Cat# 034-24024 |
| SB203580 | FUJIFILM Wako | Cat# 199-16551 |
| SB431452 | Tocris | Cat# 1614 |
| DAPI | Invitrogen | Cat# D1306 |
| Doxycycline | FUJIFILM Wako | CAS RN: 10592-13-9 |
| Penicillin | Meiji | Cat# 4987-222-63767-1 |
| Streptomycin | Meiji | Cat# 4987-222-66564-3 |
| G418 | Nacalai Tesque | CAS RN: 108321-42-2 |
| Puromycin | Nacalai Tesque | CAS RN: 58-58-2 |
| **Deposited Data** |  |  |
| RNA-seq data deposited to Gene | This paper | GEO: GSE198127 |
| **Recombinant DNA** |  |  |
| PB-EOS-C(3+)-EiP (EGFP-IRES-Puro) | Hotta et al., 2009 | N/A |
| tetracycline-inducible (Tet-ON) constitutive active form of p38 | Xu et al., 2013 | N/A |

| Critical Commercial Assays |  |  |
| --- | --- | --- |
| SYBR Green PCR Master Mix | Applied Biosystems | Cat# 4367659 |
| RNeasy Mini Kit | QIAGEN | Cat# 74104 |
| RNase-Free DNase Set (50) | QIAGEN | Cat# 79254 |
| SuperScript First-Strand Synthesis SuperMix | Invitrogen | Cat# 11752-050 |
| SuperScript™ II Reverse Transcriptase 10,000 units | Invitrogen | Cat# 18064014 |
| RNA Clean & Concentrator-5 kit | Zymo Research | Cat# R1013 |
| NEBNext® Poly(A) mRNA Magnetic Isolation Module | New England BioLabs | Cat# E7490S |
| SMART-Seq® Stranded Kit | Takara Bio | Cat# 634442 |
| Tetramethylrhodamine, Ethyl Ester, Perchlorate (TMRE) | Life Technology | Cat# T669 |
| Oligonucleotides |  |  |
| Primers for qPCR, see Table S2 | This paper | N/A |
| Software and Algorithms |  |  |
| ImageJ | ImageJ | https://imagej.nih.gov/ij/ |

## CONTACT FOR REAGENT AND RESOURCE SHARING

Further information and requests for resources and reagents should be directed to and will be fulfilled by the Lead Contact, Jun.K.Yamashita.

## EXPERIMENTAL MODEL AND SUBJECT DETAILS

### Cell lines

The human ESC line primed H1 cells were obtained from WiCell.

The human iPSC lines Ff-I14 was provided by the Center for iPS Cell Research and Application, Kyoto University.

## METHOD DETAILS

### Cell culture

hiPSC lines (FfI14) were provided by the Center for iPS Cell research and Application, Kyoto University and hESCs line (H1) was provided by WiCELL. Primed PSCs were maintained on Matrigel (Invitrogen, 1:60 dilution) coated dishes in StemFit AK02N (Ajinomoto) (Kubara et al., 2018). Cells were passaged every 3-7 days as single cells using TrypLE™ Select CTS™ (Gibco). The ROCK inhibitor Y27632 (Fujifilm, 10 μM) was added to the medium for the first 24 hr after passage.

Naïve human PSCs were cultured in the PXGL medium: Ndiff227 (Takara Bio) medium supplemented with PD0325901 (Sigma,1 μM), XAV939 (Millipore, 2 μM), Go6983 (Fujifilm, 2 μM), human LIF (Fujifilm,10 ng/ml), and penicillin/streptomycin (Meiji) on Matrigel-coated 6 well plates with mitomycin C inactivated mouse embryonic fibroblast (MEF) feeder.

AICAR (Fujifilm, 1mM) was added to the PXGL medium (Liu et al., 2019; Liu et al., 2021), and VPA (Guo et al., 2017) was added to Ndiff227 basal medium with PD0325901 (Sigma, 1 μM), human Lif (Fujifilm, 10 ng/ml), and penicillin/streptomycin) for naïve state induction. Cells were passaged on MEF feeder layers every 5-10 days as single cells using TrypLE™ Select CTS™ (Gibco). The ROCK inhibitor Y27632 (Fujifilm, 10 μM) was added to the medium for the first 24 hr after passage. The medium was changed daily.

### EOS-GFP and CA-p38 transfection

Mouse ES cells and iPS cells that are in the naïve state, and naïve state human ES cells and iPS cells have already been reported to preferentially use the distal enhancer (DE) in Oct4 transcription (Yeom et al., 1996; Takashima et al., 2014; Theunissen et al., 2014; Choi et al., 2016), while mouse Epi-SCs, human ES cells and iPS cells, known as primed state cells, preferentially use the proximal enhancer (PE) (Tesar et al., 2007; Gafni et al., 2013). PB-EOS-C(3+)-EiP (EGFP-IRES-Puro) has been described as a reporter of DE-Oct4 transcription and is used as a naïve state marker (Takashima et al., 2014; Hotta et al., 2009). To monitor the naïve to primed state conversion in living human PSC cells, we transfected PB-EOS-C(3+)-EiP (EGFP-IRES-Puro) *piggyBac* vector and pHL-EF1a-hcPBase-iC-A vector into the H1 strain of human ES cells (H1-EOS) and the human iPS cell strain Ff-I14 (Ff-I14-EOS). 24 hr after electroporation by NEPA21 (Nepagene), cells were selected by treatment of puromycin (Nacalai, 10 μg/ml) for 5 days. Transfected cells were maintained on Matrigel-coated dishes in StemFit AK02N (Ajinomoto).

For generating of CA-p38 H1-EOS cell lines, H1-EOS cells were transfected with tetracycline-inducible (Tet-ON) constitutive active form of p38 (CA-p38), which contains D176A and F327S mutated p38 cDNA in *piggyBac* (PB) vector carrying rtTA expression coupled to mCherry (Xu et al., 2013; Liu et al., 2019). mCherry positive cells after treatment of doxycycline (DOX) were purified by FACS. Transfected cells were maintained on Matrigel-coated dishes in StemFit AK02N (Ajinomoto).

pHL-EF1a-hcPBase-iC-A was used for *piggyBac* transposase, and NEPA21 (NEPA GENE) was used for electroporation.

### Flow cytometry

Cells were dissociated into single cells with TrypLE™ Select CTS™ and stained with conjugated antibodies and 4’, 6-diamidino-2-phenylindole (DAPI).

SUSD2 clone W5C5 (SUSD2-PE, Bio Legend 327406), CD75/CD75s Clone ZB55 (CD75-APC, BD 566350) and CD57 Clone NK-1 (CD57-BV421, BD 563896) were used for flow cytometry (Collier et al., 2017; Bredenkamp et al., 2019).

### Immunostaining

Immunostaining was carried out as described previously (Yamashita et al., 2000). Cells were fixed in 4% paraformaldehyde for 15 min, then blocked with Blocking One Histo (Nacalai, 1:20 dilution) / PBS+0.5% Triton for 1 hr.

Cells were stained with primary antibodies diluted in PBS+0.5% Triton and incubated overnight at 4 °C.

Secondary antibodies, anti-rabbit, mouse, and goat IgG antibodies conjugated with Alexa488 or Alexa546 (Thermo), diluted in PBS+0.5% Triton were incubated for 1 hr at room temperature.

Phosphate-buffered serine with Tween-20 was used for washing, and DAPI was used for Nuclei staining.

### Western blotting

CA-p38 H1-EOS cells were lysed in Sample buffer solution with 2-mercaptoethanol (ME) (Nacalai). The protein in whole-cell lysates were separated by using BlotTM Gel (Invitrogen) and transferred to nitrocellulose membranes.

Membranes were blocked by using Blocking One (Nacalai) for 30 min. Then membranes were incubated overnight at 4 °C with the following primary antibodies: p38 (Cell Signaling (9212S), 1:1000), phosphorylated-p38 (Thr180/Tyr182, Cell Signaling (9215S), 1:1000) and β-actin (Sigma (A5441), 1:10000) diluted in the Can Get Signal Immunoreaction Enhancer Solution Kit (Toyobo). Anti-mouse and rabbit IgG antibodies-conjugated with

Horseradish peroxidase (HRP) (Cell Signaling, 1:3000-1:1000) were also diluted in the Can Get Signal Immunoreaction Enhancer Solution Kit used for secondary antibodies.

After incubation of secondary antibodies for 2 hr at room temperature, the detection was performed using Immobilon western chemiluminescent substrate (Millipore).

### RNA isolation and quantitative reverse transcription polymerase chain reaction (RT-qPCR)

Total RNA was isolated the RNeasy Mini kit (Qiagen), and the Super-Script III (Invitrogen) was used for reverse-transcription. All qPCR reactions were performed in technical triplicate of each sample used in SYBR Green Master Mix (Applied Biosystems). Cells were normalized by endogenous control RPS18.

### Cell Metabolism Assays

Mitochondria were stained with tetramethylrhod-amine, ethyl ester (TMRE, final concentration 20nM, Life Technologies) in the relevant medium for 10 min and analyzed by confocal microscopy (Takashima et al., 2014).

### RNA-sequencing

RNA-seq was performed on primed H1-EOS, primed CA-p38 H1-EOS, AICAR induced naïve H1-EOS, CA-p38 induced naïve CA-p38 H1-EOS, and VPA induced naïve H1-EOS.

For sample preparation, primed state cells were dissociated by TrypLE™ Select CTS™, and CD75+/SUSD2+ naïve state cells were sorted in by using FACS. Total RNA was isolated by using the RNeasy Mini kit (Qiagen) and purified with the RNA Clean & Concentrator-5 kit (Zymo Research) according to the manufacturer’s instructions. Polyadenylated RNA (polyA) was enriched by using NEBNext® Poly(A) mRNA Magnetic Isolation Module (New England BioLabs). RNA-seq libraries were generated by using SMART-Seq® Stranded Kit (TaKaRa bio), and sequenced by Novaseq 6000. Reads were aligned to human genome build GRCh38.p13 and Reads Per Kilobase of transcript, per Million mapped reads (RPKM) calculation were performed by using RSEM (RNA-Seq by Expectation-Maximization).

Comparative data were downloaded from the European Nucleotide Archive (ENA) under accessions ERP006823 (Takashima et al., 2014), SRP059279 (Ji et al., 2016), SRP045911 (Sperber et al., 2015), SRP055810 (Blakeley et al., 2015), and SRP074076 (Yang et al., 2017).

All 29 fastq files are adapter trimmed by cutadapt-1.15 using trim_galore-0.4.4_dev with – stringency 3 option, and then mapped to Ensembl GRCh38 release 100 reference cDNA and ncRNA sequences using Bow-tie2 v2.2.5 with the very-sensitive-local option. Among a total of 1,800,071,774 (single or paired) reads, 1,254,839,844 (69.7%) were successfully mapped and quantified as genes with a threshold of MAPQ score ≥ 1. A total of 61,222 genes (43,813 on average) were detected for 29 samples.

The following analyses were performed using R ver. 4.0.3 after non-categorized limma voom normalization (Law et al., 2014). A total of 25 samples from naïve hESCs, primed hESCs, AICAR- and CA-p38 induced naïve hPSCs were analyzed. The heatmap (Figure 5A) was analyzed for a selected gene set of pluripotency regulators, naïve and primed state associated genes (Takashima et al., 2014, Theunissen et al., 2014, Guo et al., 2016, Guo et al., 2017, Hu et al., 2020) as depicted.

### In vitro differentiation

Naïve cells were ‘re-primed’ before in vitro differentiation. Naïve cells were passaged on Matrigel-coated dishes in StemFit AK02N (Ajinomoto) for about 1 month.

For endoderm differentiation, the medium was switched to mTeSR1 (STEMCELL Technologies) from StemFit AK02N (Ajinomoto), and cultured re-primed cells for 1 week. Then mTeSR (STEMCELL Technologies) medium was switched to RPMI1640 (Gibco)+B27 medium (RPMI1640, 2 mM L-glutamine, x1 B27) with Activin A (R&D, 100 ng/ml) and Wnt3A (Proteintech, 25 ng/ml).

The next day, the medium was changed to RPMI medium with Activin A (R&D, 100 ng/ml) and 0.2% serum.

For mesoderm differentiation, re-primed cells were cultured in StemFit AK02N (Ajinomoto) up to a confluence state, and then cells were covered with Matrigel (Invitrogen, 1:60 dilution) diluted in mouse embryonic fibroblast conditioned medium (MEF-CM) supplemented with hbFGF (Fujifilm,4 ng/ml) for 1 day. On the following day, MEF-CM was replaced with RPMI1640 (Gibco)+B27 medium (RPMI1640, 2 mM L-glutamine, x1 B27 supplement without insulin) supplemented with Activin A (R&D, 100 ng/ml) for 24 hr, followed by human Bone morphogenetic protein 4 (R&D, 10 ng/ml) and hbFGF (Fujifilm,10 ng/ml) for 4 days with no culture medium replacement. Finally, the medium was switched to RPMI1640 (Gibco)+B27 medium (RPMI1640, 2 mM L-glutamine, x1 B27), and cultured for 1 week.

For ectoderm differentiation, re-primed cells were cultured with Ndiff227 (Takara Bio) supplemented with hbFGF (Fujifilm,10 ng/ml), SB431542 (Tocris, 20 mM), and Noggin (R&D, 260 ng/ml) for 4days (Takashima et al., 2014), and then the medium was switched to Ndiff227 (Takara Bio) supplemented with hbFGF (Fujifilm,10 ng/ml) and SB431542 (Tocris, 20 mM).

### Cell growth rate determination

Naïve cells maintained on MEF feeder cells were dissociated by using TrypLE™ Select CTS™.

Dissociated cells were counted and stained with SUSD2-PE and CD75-APC. SUSD2+/CD75+ ratio detected by using TrypLE™ Select CTS™ FACS were used for calculating the number of naïve hPSCs.

## QUANTIFICATION AND STATISTICAL ANALYSIS

### Imaging analysis

Western blot data was quantified by Image J. Statistical analysis was done using a Oneway repeated analysis of variance (ANOVA), followed by the Tukey’s test as post hoc. Error bars indicate mean ± SEM.

---

**Supplementary-Figure1.**
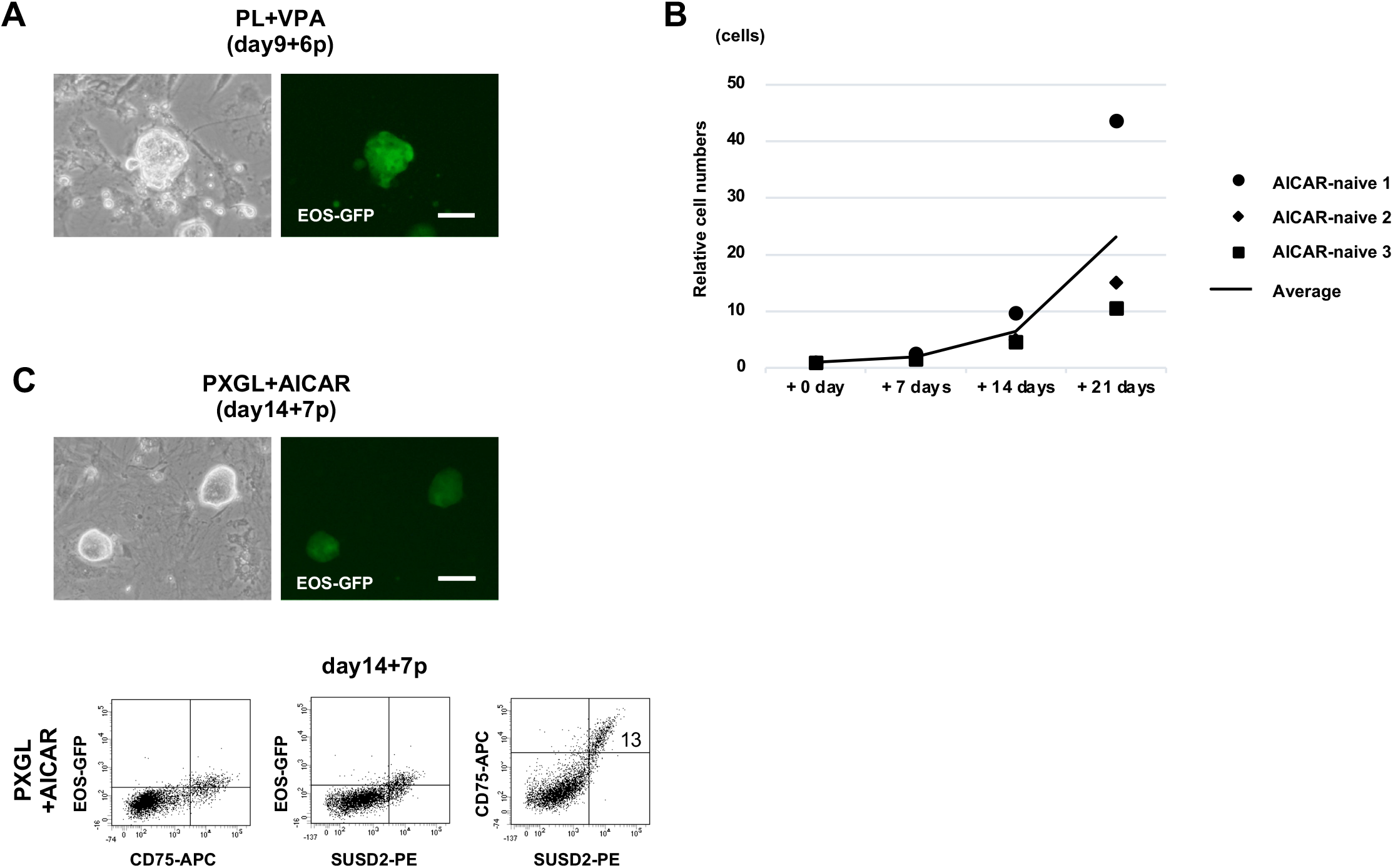
Naive conversion of human iPS cells. (A) Images of naïve hESCs (H1-EOS) induced by VPA. VPA, valproic acid (day9+6p). (B) Cell proliferation data of naive hESCs in the PXGL on MEF feeder cells. AICAR-naive 1: day14+19p, AICAR-naive 2: day14+20p, AICAR-naive 3: day14+7p at day0 in figure. (C) Images of naïve hiPSCs (Ff-I14-EOS) induced by AICAR(day14+7p) and Flow-cytometry analysis of EOS-GFP, SUSD2 and CD75 expression in naive hiPSCs (day14+7p). Scale bars: 100 μm

**Supplementary-Figure2.**
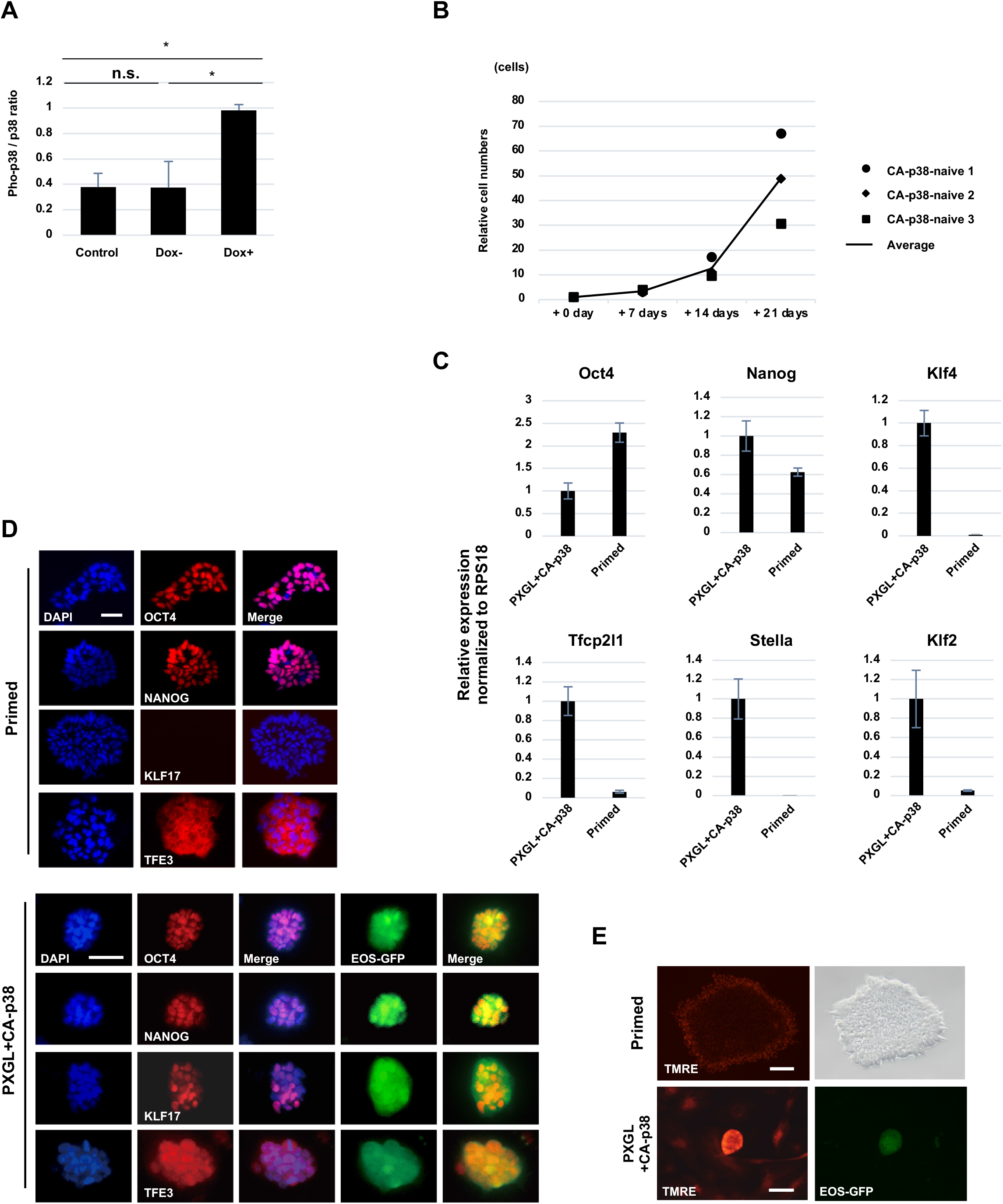
Naive reprogramming. (A) p-p38 expression analyzed by western blotting. β-Actin was used as a loading control. Error bars indicate S.E. (n=3) *p < 0.05. (B) Cell proliferation data of naive hESCs in the PXGL on MEF feeder cells. CA-p38-naive 1: day7+8p, CA-p38-naive 2: day7+16p, CA-p38-naive 3: day7+10p at day0 in figure. (C) RT-qPCR analysis of sorted SUSD2+CD75+ naive hESCs and conventional primed hESCs (CA-p38 H1-EOS). CA-p38: day7+10p. Error bars: S.D. of technical triplicates. (D) Immunostaining for OCT4, NANOG, KLF17 and TFE3 of naive conversion. CA-p38: day7+12p. (E) TMRE staining for mitochondrion dependent on mitochondrial membrane activity. Scale bars: 50 μm in D; 100 μm in E. CA-p38: day7+14p.

**Supplementary-Figure3.**
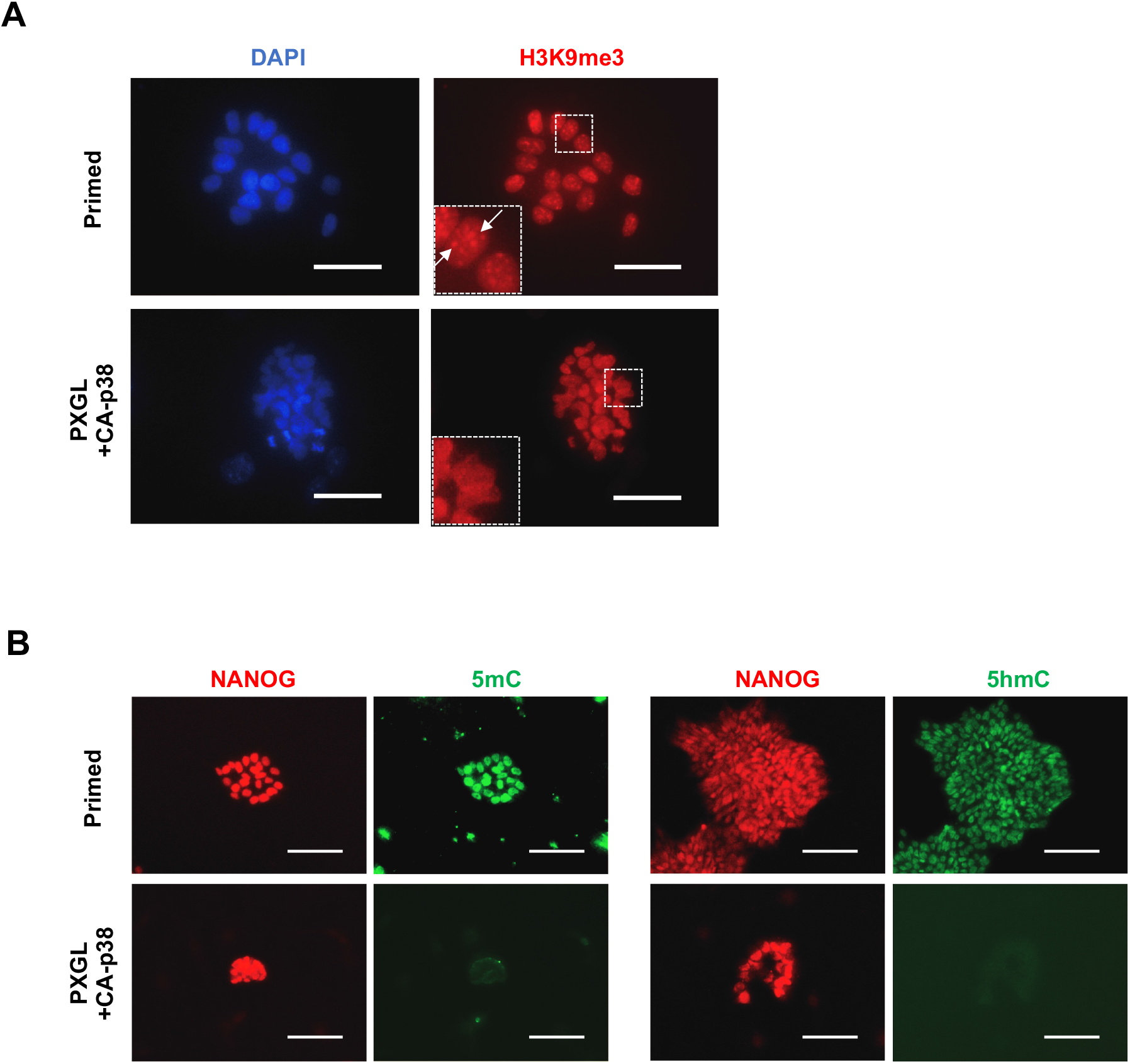
Epi-genome analysis of induced naïve hPSCs. (A) Immunostaining for H3K9me3. (B) Immunostaining for 5mC, 5hmC and NANOG. Scale bars: 50 μm in A; 100 μm in B. CA-p38: day7+12p.

**Supplementary-Figure4.**
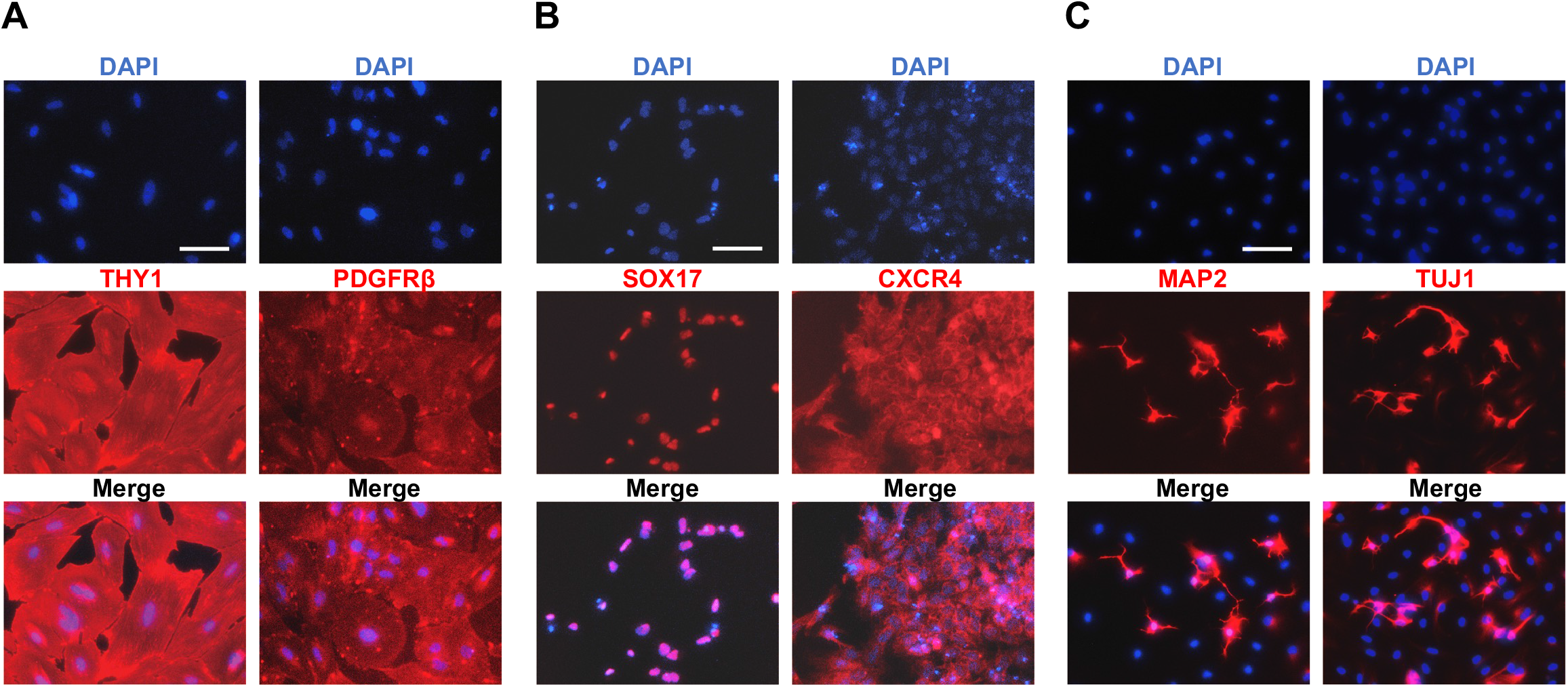
Differentiation of induced naive hPSCs. (A) Immunofluorescence for mesoderm markers THY1 and PDGFRβ. (B) Immunofluorescence for endoderm markers SOX17 and CXCR4. (C) Immunofluorescence for ectoderm markers MAP2 and TUJ1. Scale bars: 100 μm.

