## Supplementary material for "AMPK-p38 axis converts human pluripotent stem cells to naïve state": Table S1

| | | Naïve hPSC | | induced Naïve hPSC | | | | | | | | |
|---|---|---|---|---|---|---|---|---|---|---|---|---|
| | | 2016 | 2019 | 2013 | 2013 | 2014 | 2014 | 2016 | 2017 | 2019 | 2020 | 2020 |
| | | Guo et al. | Bredenkamp et al. | Gafni et al. | Chan et al. | Takashima et al. | Theunissen et al. | Qin et al. | Guo et al. | Szczerbinska et al. | Hu et al. | this study |
| | | Stem Cell Reports | Stem Cell Reports | nature | Cell stem cell | Cell | Cell stem cell | Cell reports | Development | Stem Cell Reports | SCIENCE ADVANCES | |
| | | HNES (t2iLGoY) | HNES (PXGL) | NHSM | 3iL | Reset cell | 5i/L/A | Yin-PSCs | chemically reset (cR) | FINE cell | 2iLI | |
| maintaining for naïve state related component | Medium | N2B27 medium | N2B27 medium | knockout DMEM,N2 supplement, etc. medium | TeSR1 medium | N2B27 medium | N2B27 medium | N2B27 medium | N2B27 medium | N2B27 medium | N2B27 medium | N2B27 medium |
| | Growth factors | LIF (10 ng/ml) | LIF (10 ng/ml) | LIF (20 ng/ml) | LIF (10 ng/ml) | LIF (10 ng/ml) | LIF (10 ng/ml) | LIF (10 ng/ml) | LIF (10 ng/ml) | LIF (20 ng/ml) | LIF (20 ng/ml) | LIF (10 ng/ml) |
| | ERK inhibitor | PD0325901 (1 μM) | PD0325901 (1 μM) | PD0325901 (1 μM) | PD0325901 (1 μM) | PD0325901 (1 μM) | PD0325901 (1 μM) | PD0325901 (0.5 μM) | PD0325901 (1 μM) | PD0325901 (1 μM) | PD0325901 (1 μM) | PD0325901 (1 μM) |
| | GSK3 inhibitor | CHIR99021 (1 μM) | — | CHIR99021 (3 μM) | BIO (2 μM) | CHIR99021 (1 μM) | IM-12 (1 μM) | CHIR99021 (3 μM) | CHIR99021 (0 or 0.3 μM) | — | CHIR99021 (3 μM) | — |
| | Tankyrase inhibitor | — | XAV939 (2 μM) | — | — | — | — | — | XAV939 (2 μM) | — | — | XAV939 (2 μM) |
| | ROCK inhibitor | Y-27632 (10 μM) | — | Y-27632 (5 μM) | — | — | Y-27632 (10 μM) | — | — | Y-27632 (10 μM) | — | — |
| | PKC inhibitor | Go6983 (2.5 μM) | Go6983 (2 μM) | Go6983 (5 μM) | — | Go6983 (5 μM) | — | — | Go6983 (5 μM) | — | — | Go6983 (2 μM) |
| induction for naïve state related component | Growth factors | — | — | FGF2 (8 ng/ml) | — | — | — | — | — | FGF2 (8 ng/ml) | — | — |
| | Cytokines | — | — | TGFβ1 (1 ng/ml) | — | — | Activin (20 ng/ml) | — | — | Activin (20 ng/ml) | — | — |
| | B-Raf kinase inhibitior | — | — | — | — | — | SB590885 (0.5 μM) | — | — | SB590885 (0.5 μM) | — | — |
| | Other inhibitors | — | — | JNK inhibitor (SP600125 10 μM) | — | – | Src inhibitor (WH4023 1 μM) | — | — | — | mTOR inhibitor (Torin1 10 μM) | — |
| | Transgene | — | — | — | — | Nanog<br>KLF2 | — | (YAP ※) | — | — | — | (CA-p38 ※) |
| | AMPK-p38 | — | — | p38 inhibitor<br>(SB203580 10 μM) | AMPK inhibitor<br>(Dorsomorphin 2 μM) | — | — | cAMP, AMPK activator<br>(Forskolin 10 μM) | — | — | — | AMPK activator<br>(AICAR 1mM) |
| | Others | — | — | — | — | — | — | YAP activator<br>(LPA 10 μM) | Valproic acid sodium salt (1mM) | AZD5438 (0.1 μM)<br>Dasatinib (0.1 μM) | human insulin<br>(18 μg/ml) | — |
| DNA CpG methylation | | Low | — | Low | — | Low | — | — | Low | Low | Low | Low |
| Repressive histone marks | | — | — | H3K27me3: low | H3K27me3: low | H3K9me3: low<br>H3K27me3: low | H3K27me3: low | H3K9me3: low | — | H3K9me3: low | H3K27me3: low | H3K9me3: low |
| Predominant OCT4 enhancer | | — | — | Distal | — | Distal | Distal | — | Distal | — | Distal | Distal |
| cell surcafece marker | | — | CD7, CD75, CD77, CD130, SUSD2 | E-CAD | — | — | — | — | — | CD75, CD130 | — | CD75, SUSD2 |
| transcriptome | | KLF4, TFCP2L1, KLF17, STELLA, DPPA5, KLF2, TBX3 | KLF17, TFCP2L1, KLF4, DPPA5 | — | High: GP130, KLF4, TBX3, STELLA, KLF5 | High: TBX3, REX1, STELLA, TFCP2L1, KLF2, KLF4, GBX2, ESRRB | High: KLF2, KLF4, REX1, STELLA, DPPA5, TFCP2L1 | High: GP130, TBX3, TFCP2L1, STELLA, HERVH-Gap, HERVH-Pol | High: KLF17, TFCP2L1, STELLA, KLF4, TBX3 | High: KLF4, KLF5, KLF17, STELLA, LTR7Y, TBX3 | High: KLF4, KLF2 | High: KLF2, KLF4, KLF17, TFCP2L1, STELLA |
| TFE3 nuclear localization | | — | — | YES | — | YES | — | — | — | YES | YES | YES |
| Mitochondrial membrane activity and depolarization | | High | — | — | — | High | — | — | — | — | High | High |
