## Supplementary material for "AMPK-p38 axis converts human pluripotent stem cells to naïve state": Table S2

Table S2. Primer Sequences for qPCR, Related to STAR Methods, Figure 1 and S2

| Gene | Forward primer 5'-3' | Reverse primer 5'-3' |
|---|---|---|
| RPS18 | ACTCAACACGGGAAACCTCA | AACCAGACAAATCGCTCCAC |
| OCT3/4 | TGTACTCCTCGGTCCCTTTC | TCCAGGTTTTCTTTCCCTAGC |
| NANOG | CAGTCTGGACACTGGCTGAA | CTCGCTGATTAGGCTCCAAC |
| KLF4 | GATGGGGTCTGTGACTGGAT | CCCCCAACTCACGGATATAA |
| TFCP2L1 | CTCAGGTGCTGACTTGCTGA | ATGGCGTGGTACACAGACAG |
| STELLA | TCTCCACAAATGCTCACCGA | TCTTCTTTCATGCGTACGAACTCC |
| KLF2 | CATCTGAAGGCGCATCTG | CGTGTGCTTTCGGTAGTGG |
